# Whole-mount acetylcholinesterase (AChE) staining reveals unique motor innervation of the lamprey oral region: with special reference to the evolutionary origin of the vertebrate jaw

**DOI:** 10.1101/2025.04.10.648078

**Authors:** Motoki Tamura, Daichi G. Suzuki

## Abstract

The evolutionary change of the trigeminal nerve-innervation pattern is essential to reveal the mechanism underlying jaw acquisition. However, the homology of the branches between gnathostomes (jawed vertebrates) and cyclostomes (living jawless vertebrates) remains unclear. In this study, we focused on a sub-branch called the ramus subpharyngeus, which belongs to the second branch of the lamprey trigeminal nerve and projects to the lower lip, investigating whether it contains motor components. To visualize motor fibers, we performed acetylcholinesterase (AChE) staining, a histochemical method that visualizes intrinsic activities of the catabolic enzymes produced by motor neurons and muscle fibers. As a result, we found AChE staining signals that correspond to the innervation course of the ramus subpharyngeus. To confirm that the signals are not produced by the motoneuronal somata nor muscle fibers, we conducted gene expression analysis by *in situ* hybridization. The results support that the signals mark the motor fibers. Based on these results, we propose that the lamprey oral apparatus is chiefly controlled by the second (i.e., pre-mandibular) branch of the trigeminal nerve and further suggest that a drastic reorganization of the anterior craniofacial region occurred during the acquisition of the vertebrate jaw.

## Introduction

The acquisition of the jaw is a pivotal event in the evolutionary history of vertebrates (Forey and Janvier, 1993). Still, it remains unclear how this effective predatory apparatus could evolve from the ancestral jawless fishes (Kuratani, 2005; Miyashita, 2016; Romer and Parsons, 1986). As this organ is controlled by the trigeminal nerve (i.e., the fifth cranial nerve, V), the evolutionary modification of its innervation pattern is a key to solving this issue.

The trigeminal nerve generally shows three main branches or rami (hereafter, a ramus is abbreviated as r). Among these branches, the second (the maximally nerve, rV_2_) and the third (the mandibular nerve, rV_3_) ones innervate the upper and lower jaw, respectively (Goodrich, 1930; Mallatt, 1996; Romer and Parsons, 1986). Notably, only the rV_3_ possesses motor components in jawed vertebrates (with an exception of holocephalans), controlling their jaw movements (Mallatt, 1996).

The cyclostomes are the sole surviving jawless vertebrates, including lampreys and hagfish (Kuraku and Kuratani, 2006; Kuraku et al., 1999). In contrast to the jawed vertebrates, their trigeminal nerve contains motor components in both rV_2_ and rV_3_ (Hardisty and Potter, 1971; Johnston, 1905; Kuratani et al., 2004; Lindström, 1949). This difference in the motor innervation patterns between these two clades may thus shed light on the origin of the jaw. Nevertheless, it involves a long-standing problem on the homology of the trigeminal nerve branches between them to answer this question.

Classically, the rV_2_ and rV_3_ of the jawed vertebrates have been thought to be homologous to the cyclostome second and third branches, respectively (Johnston, 1905; Mallatt, 1996, 2008). However, recent morphological studies have cast doubt on this homology, and thus the second and third branches of the lamprey are now denoted as the rV_2/3A_ and rV_2/3B_, respectively (Oisi et al., 2013; Tamura et al., 2023; Yokoyama et al., 2021). In particular, the lower lip of the lamprey is generally believed to be innervated by the rV_2/3B_ (Johnston, 1905; Richardson et al., 2010). However, this region is in fact supplied not only by the rV_2/3B_ but also by a sub-branch of the rV2/3_A_ called the r. subpharyngeus (rV_2/3Asubp_; Lindström 1949; see also Yokoyama et al. 2021). It has been controversial whether the rV2/3_Asubp_ contains motor component (Johnston, 1905; Richardson et al., 2010) or not (Alcock, 1898; Gaskell, 1900), being critical to clarify the homology of the trigeminal nerve branches between jawed vertebrates and cyclostomes and thus to understand evolutionary origin of the vertebrate jaw.

Several methods are available to identify motor components in peripheral nerves, but most of them are hardly applicable for the lamprey trigeminal nerve. First, there is no good antibody against motor neuron markers for the lamprey. Second, the complex lifecycle of this animal prevents the establishment of genetic lines. Although I-SceI meganuclease-mediated tissue-specific transient transgenesis and CRISPR-Cas9-mediated knock-in techniques are reported for the lamprey, only mosaic F_0_ animals can be obtained (Hockman et al., 2019; Parker et al., 2014; Suzuki et al., 2021). Last, as the rV_2/3A_ and rV_2/3B_ are proximally juxtaposed in the lower lip, it is technically difficult to microinject neuronal tracers to label these branches separately.

In this study, we therefore employ the acetylcholinesterase (AChE) staining method, which chemically detects intrinsic AChE activity by *in situ* precipitation of copper ferrocyanide (Karnovsky and Roots 1964). Although a few previous studies have reported whole-mount application for invertebrates (Bery and Martínez, 2011; Denker et al., 2008), this technique is principally used for sectioned materials (Karnovsky and Roots, 1964; Kusunoki, 1981; Kusunoki et al., 1973; Kusunoki et al., 1982) and no whole-mount analysis on vertebrate embryos has been reported.

Here, we first report the whole-mount AChE staining results for the lamprey embryos and prolarvae, with special reference to their oral region. We found AChE signals in the innervation course of the rV_2/3Asubp_. As AChE signals are found not only in motor fibers but also in neuronal somata and muscles, we then conducted in situ hybridization analysis for *AChE* and *Muscle Actin 2* (*MA2*) genes. As a result, we confirmed that the AChE signals in the innervation course of the rV_2/3Asubp_ were not produced by motoneuronal somata or muscles, suggesting that rV_2/3Asubp_ contains motor fibers. Based on these results, we discuss the relationship between oral structure and innervation trigeminal motor components of lampreys in light of the evolutionary origin of the vertebrate jaw.

## Materials & Methods

### Ethics Statement

All procedures in this study were performed in compliance with the guidelines for animal use of the Animal Care Committees at University of Tsukuba (specific approval is not required for experimentation on fishes under the Japanese law, Act on Welfare and Management of Animals). During the investigation, every effort was made to minimize suffering and to reduce the number of animals used.

### Animals

Adult male and female lampreys (*Lethenteron camtschaticum*) were collected in the Shiribeshi-Toshibetsu River, Hokkaido, Japan, during the breeding season (late May to June) from 2023 to 2024. They were brought into the laboratory and kept in aquaria with an enriched environment where water was aerated and filtered continuously. Sexually mature lampreys were deeply anesthetized in 0.1% tricaine methanesulfonate (MS-222, Sigma, E10521) for artificial fertilization; mature eggs were collected from females and fertilized in vitro by sperm, and then kept in 1×Steinberg solution at 8–12 °C. Embryos and prolarvae were staged morphologically based on the method of Tahara (1988). They were fixed with 4% paraformaldehyde in 0.1 M phosphate-buffered saline (PFA/PBS) at stages (St.) 24–28. After fixation with 4% PFA/PBS at room temperature (RT) for 12 hours, embryos were dehydrated in a graded methanol series (25%, 50%, 75% and 100%), and stored at −20 °C.

### AChE staining

Embryos were fixed with 4% PFA/PBS at 4 °C for 10 minutes. Fixed specimens were treated with 7.14% MgCl_2_ for 5 minutes and washed three times for 5 minutes each in PBT (0.1% Tween 20 in PBS). Then, specimens were incubated in Karnovsky’s solution at RT as described by Bery and Martínez (2011). Stained specimens were post-fixed in 4% PFA/PBS at 4 °C overnight. After post-fixation, specimens were washed in PBT, clarified with LUCID (Mizutani et al., 2018) and then examined under a biological microscope (Nikon, NI-FLT6) and photographed by a CCD camera (Nikon, DS-Ri1).

### Isolation of cDNA clones of lamprey

We performed tBLASTN (v2.9.0+; Camacho et al., 2009) search for *AChE* in a gene model GRAS-LJ (Kadota et al., 2017) for the *L. camtschaticum* genome assembly LetJap1.0 (BioProject: PRJNA192554) using the amino acid sequence of the human AChE (GenBank: NP_000656.1). The resulting transcripts were used as queries for searches in global protein databases at NCBI and only one resulting *acetylcholinesterase* hit was retained. To perform specific *in situ* hybridization experiments, we subcloned by polymerase chain reaction (PCR) using specific primers added SP6 promoter sequences to 3’ region (*AChE* Forward: 5’-GGACGTCTACGATGGCAAGTACCTGGCCTA-3’; Reverse:

5’-ATTAACCCTCACTAAAGGGAACTCGATCTCGTAGCCGTGCA-3’). PCR fragments were checked by agarose gel electrophoresis for specificity, and the rest of the PCR reaction purified using Exo-SAP-IT *Express* solution (Thermo Fisher, 75001.200.UL) and then sequenced.

### Phylogenetic tree analysis of Lamprey *AChE*

Amino acid sequences were aligned using the FFT-NS-2 strategy from MAFFT v.7 and trimmed by trimAl. A phylogenetic tree was produced by the maximum likelihood method using IQ-TREE multicore version 1.6.12 (Nguyen et al., 2015), using the LG+G4 model. A total of 1,000 bootstrap replicates were used to assess node confidence.

### Whole-mount *in situ* hybridization

Whole-mount *in situ* hybridization was performed according to the method described by Murakami et al. (2001).

Antisense RNA probes were transcribed using T3 RNA polymerase (Roche, 11031163001) in conjunction with digoxigenin conjugated dUTPs (Roche, 11277073910) following standard protocols. Specimens were treated with a mixture of hydrogen peroxide and methanol (1:5) overnight for bleaching, and were rehydrated in PBT. The samples were digested with 10 mg/ml proteinase K (Invitrogen, AM2546). They were post-fixed for 20 minutes with 4% PFA/PBT containing 0.2% glutaraldehyde, then washed with PBT, and prehybridized in hybridization buffer (50% formamide, 5× SSC, 1% SDS, 1× Denhardt’s Solution (Nacalai, 10727-74), 50 µg/ml heparin sulfate, 5 mM EDTA, 0.1% CHAPS) for 90 minutes at 70 °C. The specimens were then incubated in a hybridization buffer with 0.1 mg/ml DIG-labeled RNA probe (Roche, 11277073910) for 48 hours at 70 °C. After hybridization, the specimens were washed twice in 50% formamide, 5× SSC, and 1% SDS for 30 minutes at 70 °C, and the solution was substituted gradually with 10 mM Tris-HCl (pH 7.5) containing 0.5 M NaCl and 0.1% Tween 20 (TBST). RNaseA was added to a final concentration of 0.05 mg/ml and the specimens were incubated for 30 minutes at RT. The samples were washed twice with 2× SSC in 50% formamide for 30 minutes at 70 °C, twice in 2× SSC containing 0.3% CHAPS for 30 minutes at 70 °C, and twice in 0.2× SSC containing 0.3% CHAPS for 30 minutes at 70 °C. For immunological detection, the embryos were blocked with TBST containing 0.5% blocking reagent (Roche, 11277073910) for 90 minutes, and incubated with alkaline phosphatase (AP)-conjugated anti-digoxigenin Fab fragments (diluted 1:4000; Roche 11093274910), at 4 °C overnight. The specimens were washed ten times for 30 minutes each in TBST at RT. Alkaline phosphatase activity was detected with 20 µl/ml NBT/BCIP Stock Solution (Roche, 11681451001) in NTMT (100 mM Tris HCl pH 9.8, 100 mM NaCl). Stained specimens were fixed in 4% PFA/PBS.

### Cryosectioning

The specimens which were performed *in situ* hybridization were immersed in a graded series of sucrose (10%, 30%). Then, samples were embedded in Tissue-Tec O.C.T. Compound (Sakura Finetek, Japan), and stored at −80 °C. Frozen sections (16 µm) were prepared using a cryostat (Leica CM1860). After washing out the O.C.T. Compound, the sections were mounted with ProLong Diamond Antifade Mountant (Invitrogen, P36961).

### Whole-mount immunofluorescence

Immunofluorescence with anti-acetylated tubulin monoclonal antibody (Sigma, T6793) was performed according to the method described by Kuratani et al. (1997) with some minor modifications as described below. The samples were soaked in a 10:1 mixture of 30% hydrogen peroxide solution and 100% methanol and put under a fluorescent light for 12 hours at RT for bleaching. After 12 hours, embryos were washed in TBST containing 5% dimethyl sulfoxide (TSTd) for 3 hours at RT. After washing, the samples were sequentially blocked with 5% non-fat dried milk in TSTd (TSTM). This was followed by incubation in the primary antibody (1:1000 in TSTM) and DAPI (D9564, 1 mg/mL; Sigma-Aldrich) for 3 days at RT. After washing with TSTd, samples were incubated with secondary antibody (life technologies, Alexa fluor 488, A-21422) diluted 1:500 in TSTM for 2 days. For counterstaining, YOYO-1 equivalent solution (Abcam, ab275546) was used. After a final wash in TSTd, the embryos were clarified with LUCID (Mizutani et al., 2018), and then examined under a microscope (Nikon, NI-FLT6) and photographed by a CCD camera (Nikon, DS-Ri1).

### *In situ* hybridization combined with immunofluorescence

Whole-mount *in situ* hybridization was performed as described above. Subsequently, the samples were washed several times with TSTd. Immunofluorescence with anti-acetylated tubulin monoclonal antibody were performed as described above. After cryosectioning and mounting, the samples were examined under a fluorescence microscope (Nikon, NI-FLT6) with a light-emitting diode (LED) illuminator (Nikon, D-LEDI-C) and photographed by a CCD camera (Nikon, DS-Ri1).

## Results

### AChE staining

To investigate the motor innervation in the lamprey embryos and prolarvae, we first performed whole-mount acetylcholinesterase (AChE) staining (Figures 1, 2 and S1). In the Arctic lamprey *L. camtschaticum*, cranial nerves appear from St. 24 (Kuratani et al., 1997). Accordingly, we observed embryos from St. 24 onwards.

**Fig. 1.**
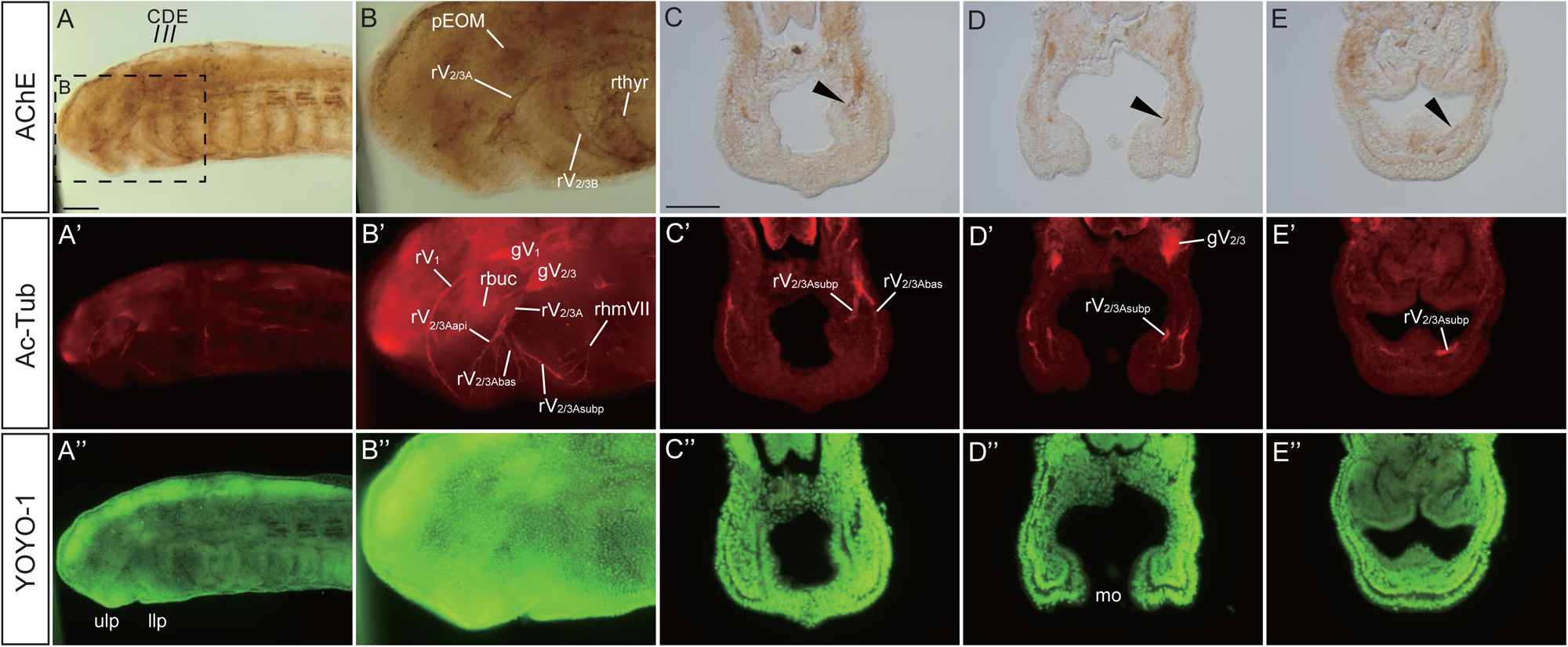
Triple whole-mount AChE staining, immunofluorescence, and YOYO-1 staining for an Arctic lamprey (*L. camtchaticum*) prolava at St. 26. (A–E) Whole-mount AChE staining images. (A’–E’) Immunofluorescence images with anti-acetylated tubulin antibody (Ac-Tub). (A’’–E’’) YOYO-1 images for visualizing the cell nuclei. (B–B’’) Enlarged image of the area enclosed by the dashed line in A–A’’. (C–E, C’–E’, C’’–E’’) Transversal section at the upper lip (C–C’’), the mouth opening (D–D’’) and lower lip level (E–E’’). Arrowheads indicates the rV_2/3Asubp_. Abbreviations: gV_1_, ophthalmic ganglion; gV_2/3_, trigeminal ganglion of V_2/3_; llp, lower lip; mo, mouth opening; pEOM, extra-ocular muscle primordium; ph, pharynx; rbuc, buccal branch of facial nerve; rhm, hyomandibular branch of facial nerve; rV_1_, ophthalmic branch of trigeminal nerve; rV_2/3A_, second branch of trigeminal nerve; rV_2/3Aapi_, apical branch of V_2/3A_; rV_2/3Abas_, basilar branch of V_2/3A_; rV_2/3Asubp_, subpharyngeal branch of V_2/3A_; rV_2/3B_, third branch of trigeminal nerve; ulp, upper lip; vel, velum. Scale bars: 100 µm for (A, applied for A–A’’) and 50 µm for (C, applied for C–E, C’–E’, C’’–E’’).

At St. 24, AChE signals were detected in the somite (Fig. S1A), but not in the head region. Similar patterns were observed at St. 25, with AChE signals evident in myotomes but absent in the head region (Fig. S1B).

At St. 26, AChE signals appeared in the head region (Fig. 1). The fibers of the cranial nerves were detected in the rV_2/3A_, rV_2/3B_, the facial nerve (rhmVII), the glossopharyngeal nerve, and the vagus nerve (Fig. 1A, A’, B and B’). The ophthalmic nerve (rV_1_) was not observed, which is consistent with the absence of the motor component in this branch. Cross-sections showed that the rV_2/3A_ with AChE staining signals was divided into two sub-branches at upper lip level, and the inner one (i.e., the rV_2/3Asubp_) passed toward buccal eminence (Fig. 1C–E, arrowheads). Then, the rV_2/3Asubp_ entered the ventral pharyngeal part below the velum. This part is the primordium of the ventromedial longitudinal bar primordium (pVMLB), which develops into the piston cartilage in adult lampreys (Armstrong et al., 1987; Hardisty and Potter, 1971; Johnels, 1948; Rose and Reiss, 1993). The AChE signals were also detected in muscle precursors. In particular, a prominent signal around the optic vesicle (Fig. 1B) indicated the presence of extraocular muscle primordium (Suzuki et al., 2016).

By St. 27, the cranial nerves of lampreys develop extensively (Kuratani et al., 1997). AChE signals were observed along the courses of the cranial nerves including the rV_2/3A_ and rV_2/3B_ (Fig. 2A, A’). At the velar level, the rV_2/3Asubp_ signals were found in the pVMLB (Fig. 2B–D, black arrowheads). At the same time, dense signals were found under the pVMLB, presumably corresponding to the lower lip muscles innervated by the rV_2/3Asubp_ (white allowheads). Nevertheless, the rV_2/3Asubp_- and lower lip muscle-signals were distantly located at this level in cross sections. After St. 29, the AChE signals of the cranial nerves were masked by strong signals of muscles (not shown).

**Fig. 2.**
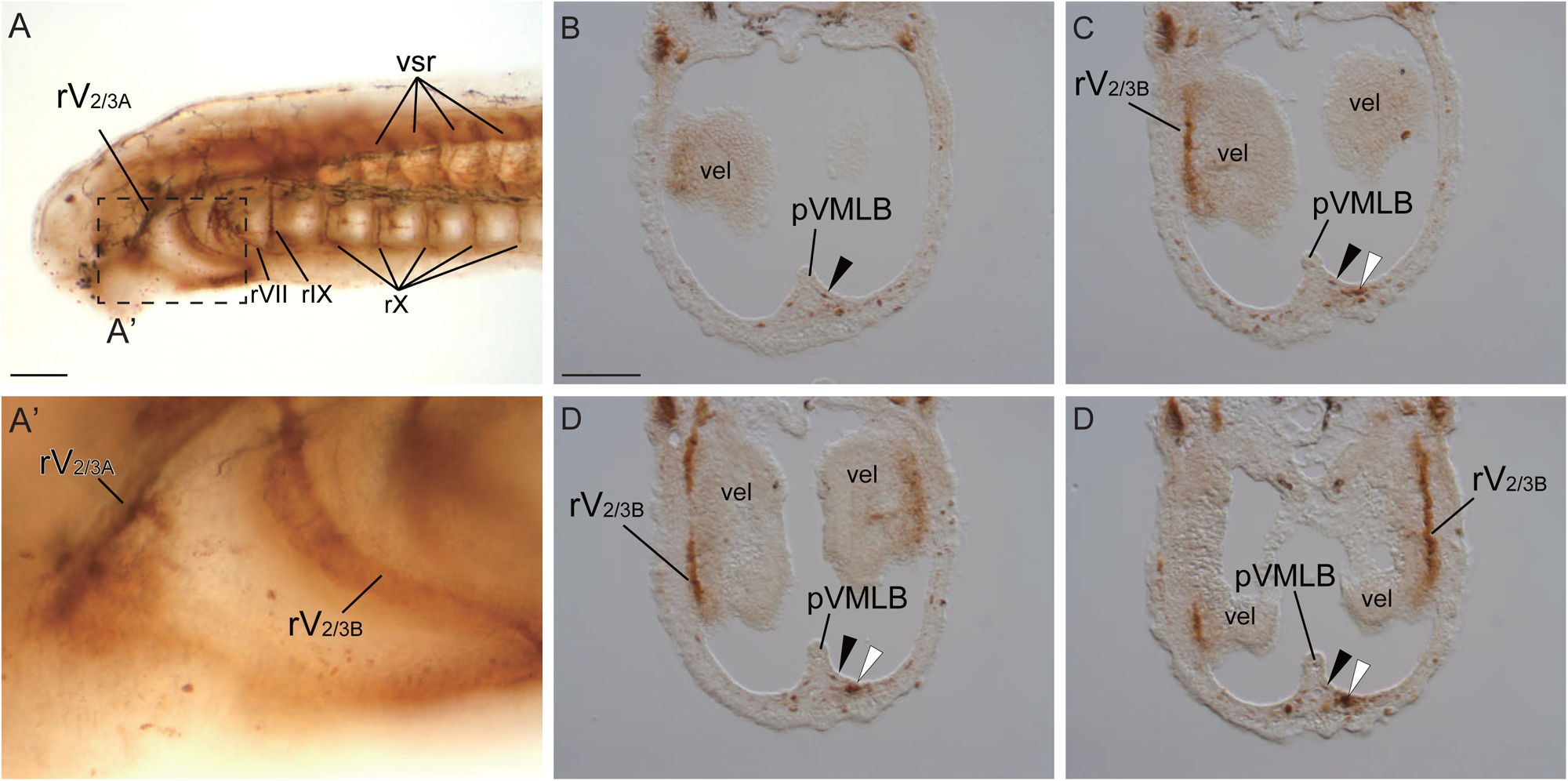
Whole-mount AChE staining for a prolava at St. 27. (A) Whole-mount AChE staining image. (A’) Enlarged image of the area enclosed by the dashed line in A. (B–D) Serial transversal sections of stained specimen of AChE staining at the velar level. Abbreviations: pVMLB, ventromedial longitudinal bar primordium; rV_2/3A_, second branch of trigeminal nerve; rV_2/3B_, third branch of trigeminal nerve; rVII, facial nerve; rIX, glossopharyngeal nerve; rX, vagus nerve; vel, velum; vsr, ventral root of spinal nerve. Scale bars: 100 µm for (A) and 50 µm for (B, applied for B–D).

Although the AChE signal in the pVMLB appears to be the motor fibers, there remains a possibility that the signal priginates from motoneuronal somata or muscles. To address this issue, we next conducted gene expression analyses as follows.

### Identification and phylogenetic analysis of the lamprey AChE gene

To determine the position of cholinergic neuronal somata and muscles, we focused on two genes: *AChE* and *MA2*. The latter was isolated in a previous study (Kusakabe et al., 2004), but the former has not been isolated so far. Thus, we first surveyed a previously published gene model for *L. camtschaticum* genome assembly (Kadota et al., 2017) and identified an *AChE* gene candidate. Previous studies have shown that gnathostomes possess two types of cholineesterases (Pezzementi et al., 2011): AChE and butyrylcholinesterase (BChE). Our phylogenetic analysis indicated that the candidate gene was an orthologue of gnathostome *AChE* (Fig. 3). Based on this result, we next investigated the expression pattern of lamprey *AChE* and *MA2*.

**Fig. 3.**
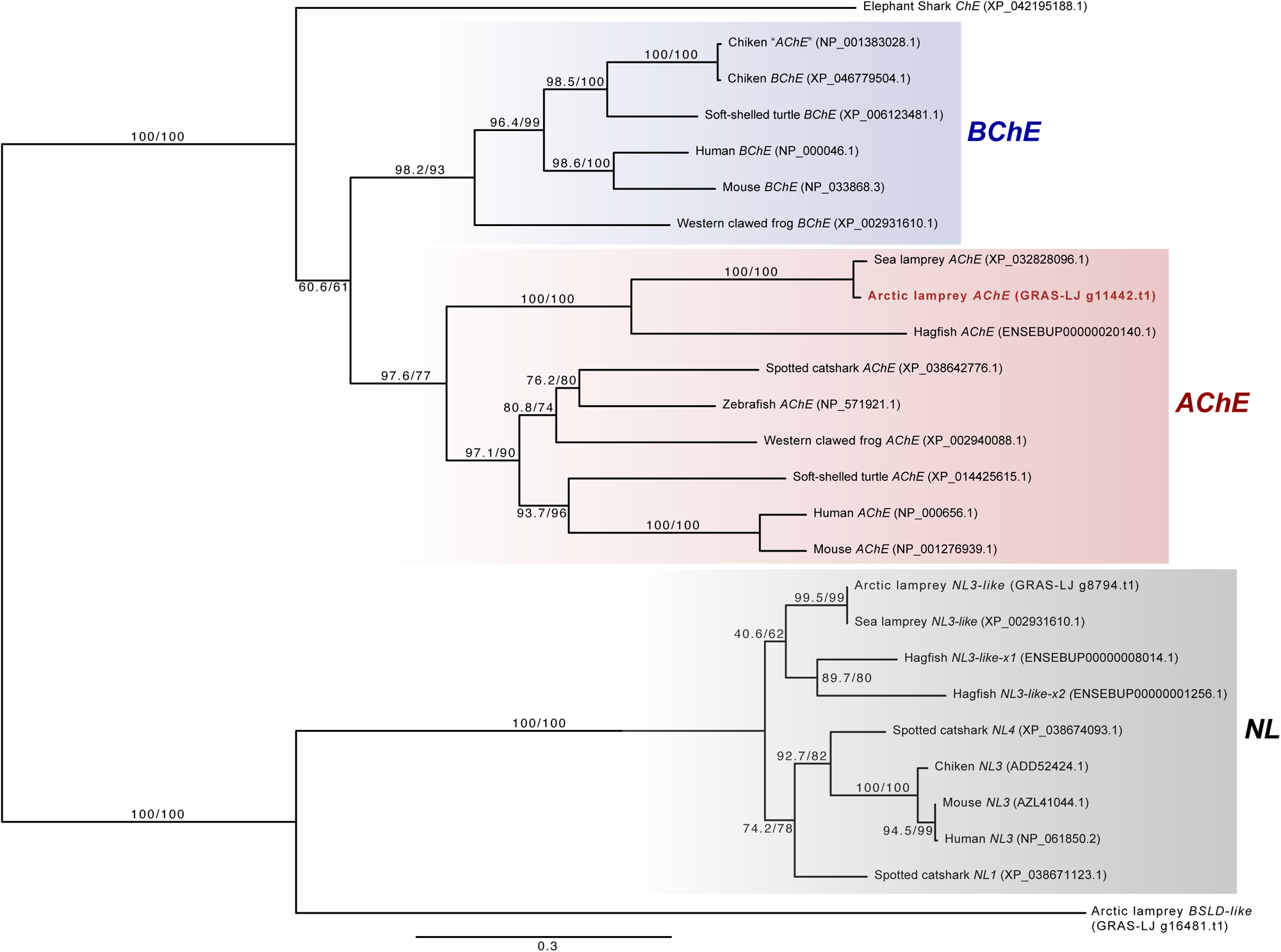
Molecular phylogenetic tree for *AChE* proteins. For Arctic lamprey genes, the sequence IDs in GRAS-LJ are indicated. Except for this species, accession numbers or Ensembl IDs are shown for respective species. The tree was constructed using the ML method. The numbers at the nodes represent bootstrap values.

### Expression pattern of *AChE* and *MA2*

To determine the position of acetylcholinergic neuronal somata and muscles in prolarval lampreys, we performed *in situ* hybridization analysis of *AChE* and *MA2* genes. For this analysis, we used St. 27 prolarvae because the signals of the AChE staining were well detected in the head region at this stage.

The lamprey *AChE* was expressed in the motor nuclei of the cranial nerves, the primordium of the extraocular muscles, the upper and lower lip muscles, the velar muscle, and the pharyngeal muscles at St. 27 (Fig. 4). While the AChE staining signal was detected in myotomes, *AChE*-positive cells were not found in them. Notably, the cross-section at the velar level showed that *AChE*-positive cells were not present in the pVMLB. These results suggest that the AChE-staining signal at the ventral pharyngeal part was not produced by cholinergic neuronal somata.

**Fig. 4.**
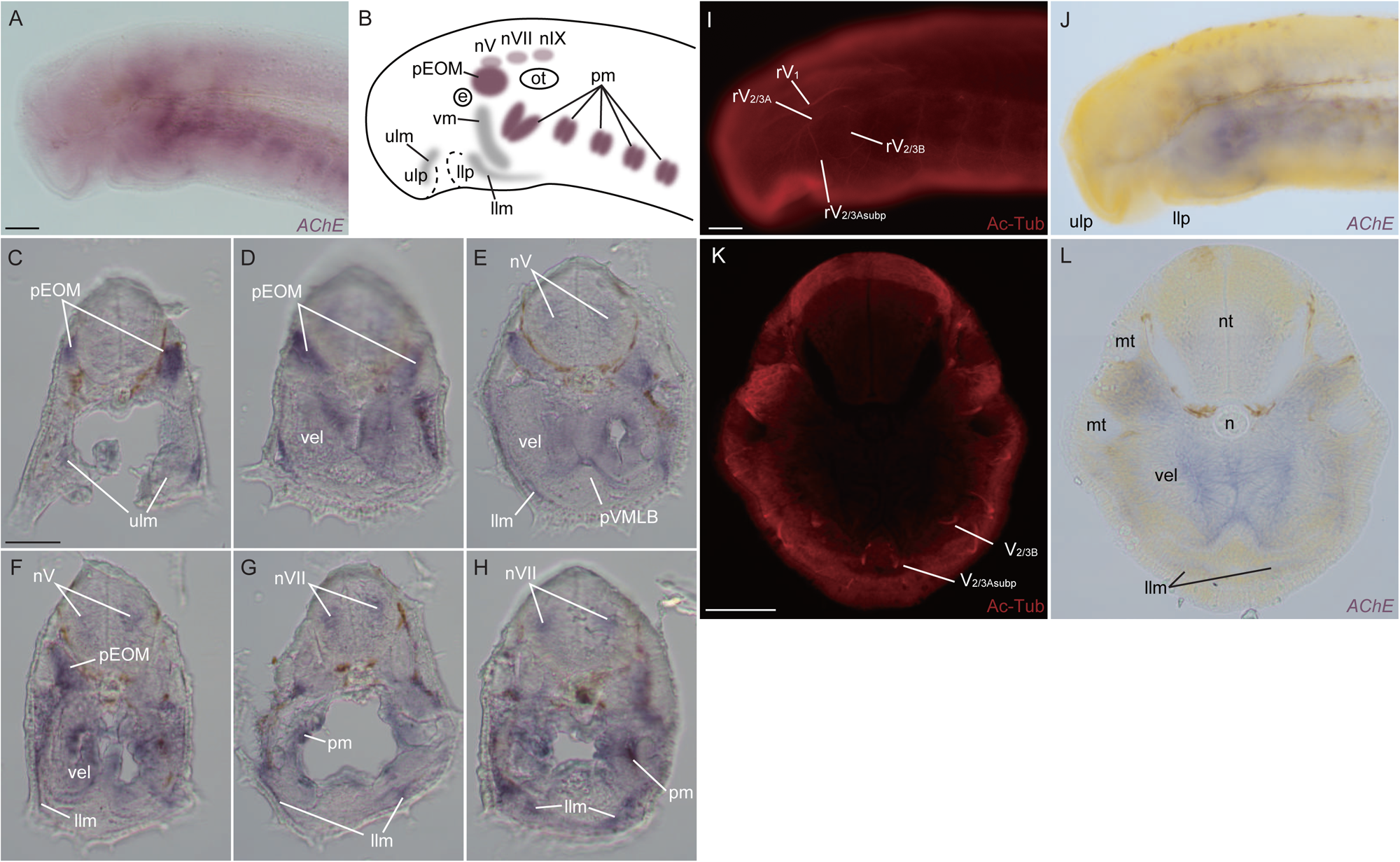
Whole-mount *in situ* hybridization of *AChE* in lamprey prolava at St. 27 (A) Lateral view. (B) Schematic diagram showing the expression pattern in A. (C–H) Transversal sections at the mouth opening-(C), the velum-(D–F) and the pharynx-levels (G, H). (I, J) Double *in situ* hybridization and immunofluorescence for *AChE* and acetylated tubulin, respectively. (K, L) Transversal section at the velum level. Abbreviations: e, eyeball; pEOM, extra-ocular muscle primordium; pm, pharyngeal muscles; pVMLB, ventromedial longitudinal bar primordium; llm, lower lip muscles; llp, lower lip; mo, mouth opening; mt, myotome; n, notochord; nV, trigeminal motor nucleus; nVII, facial motor nucleus; nIX, glossopharyngeal motor nucleus; nt, neural tube; ot, otic capsule; pEOM, extra-ocular muscle primordium; ph, pharynx; rV_1_, ophthalmic branch of trigeminal nerve; rV_2/3A_, second branch of trigeminal nerve; rV_2/3Asubp_, subpharyngeal branch of V_2/3A_; rV_2/3B_, third branch of trigeminal nerve; ulm, upper lip muscle; ulp, upper lip; vel, velum; vm, velar muscle. Scale bars: 100 µm for (A, applied also for B; I, applied also for J) and 50 µm for (C, applied for C–H; K, applied also for L).

The reason why *AChE*-positive cells were not found in myotomes may be that our RNA-probe templates were synthesized from adult lamprey cDNA library; Atkins and Pezzementi (1993) reported that the different AChE isoforms are produced in the skeletal muscles between ammocoetes and adult Sea lamprey (*Petromyzon marinus*). If it is also the case for our animal (*L. camtschaticum*), the difference in mature *AChE* mRNA sequences might prevent signal detection in prolarval myotomes by *in situ* hybridization.

The lamprey *MA2* was expressed in the muscles of the upper and lower lips, the velum, the myotomes, and the pharyngeal arches at St. 27 (Fig. 5), consistent with previous reports (Kariyayama et al., 2023; Kusakabe et al., 2004; McCauley and Bronner-Fraser, 2006; Yokoyama et al., 2021). Anterior to the otic region, the axial mesoderm was divided into two parts: the supraoptic and infraoptic muscles. The *MA2*-positive cells were not found in the extraocular muscle primordium, as these cells had not yet differentiated into muscles (Suzuki et al., 2016). Cross-sections revealed *MA2*-positive cells in the body-wall muscles at the velar level. However, no *MA2*-signals were found in the pVMLB. These results suggest that the AChE staining signal in pVMLB is not produced by muscles either.

**Fig. 5.**
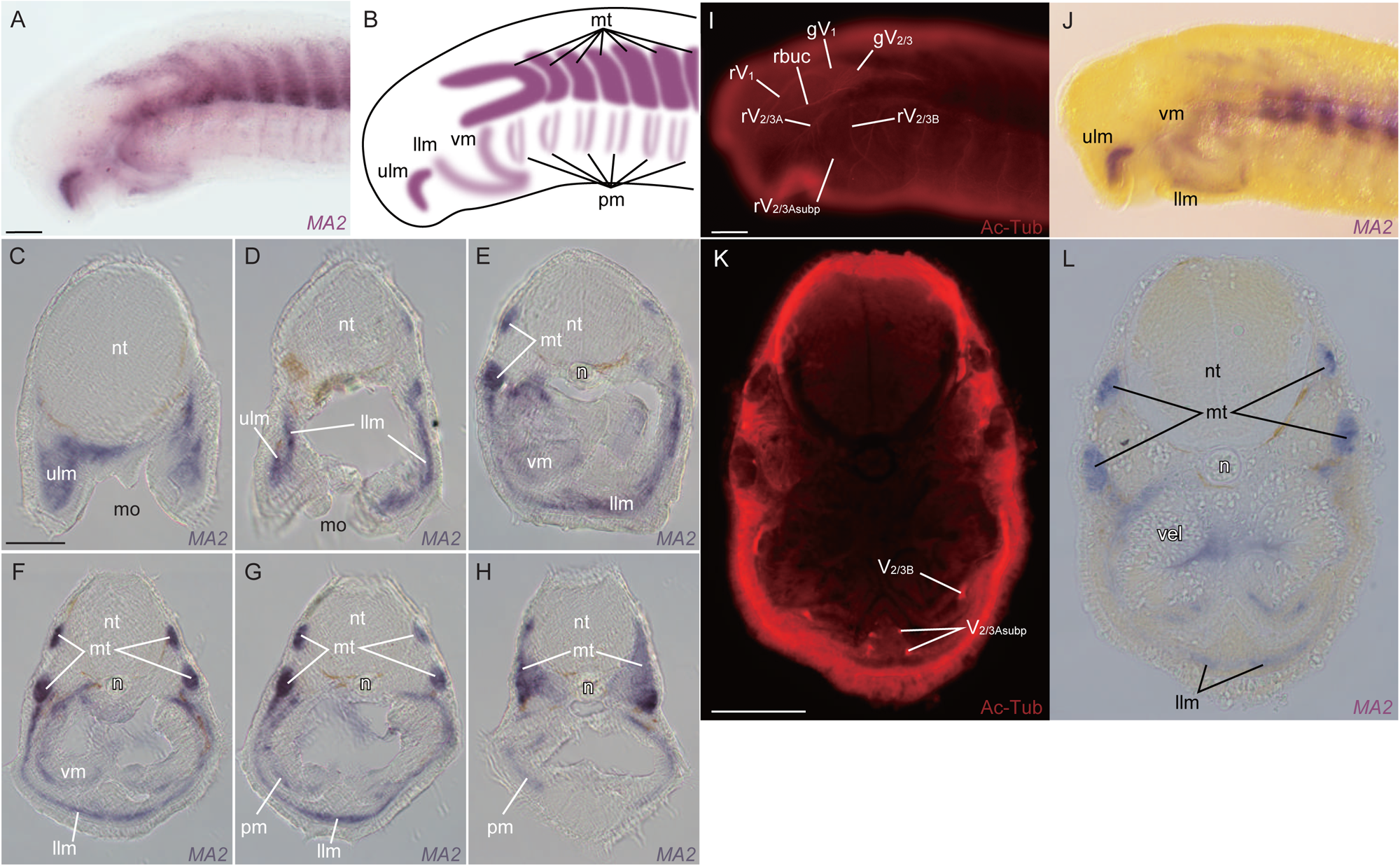
Whole-mount *in situ* hybridization of *MA2* in lamprey prolava at St. 27. (A) Lateral view. (B) Schematic diagram showing the expression pattern in A. (C–H) Transversal section at the mouth opening-(C, D), the velum-(E–G) and the pharynx-levels (H). (I, J) Double *in situ* hybridization and immunofluorescence for *MA2* and acetylated tubulin, respectively. (K, L) Transversal section at the velum level. Abbreviations: gV_1_, ophthalmic ganglion; gV_2/3_, trigeminal ganglion of rV_2/3_; llm, lower lip muscle; llp, lower lip; mo, mouth opening; mt, myotome; n, notochord; n. V, trigeminal motor nucleus; n. VII, facial motor nucleus; n. IX, glossopharyngeal motor nucleus; nt, neural tube; ot, otic capsule; pm, pharyngeal muscles; rV_1_, ophthalmic branch of trigeminal nerve; rV_2/3A_, second branch of trigeminal nerve; rV_2/3Asubp_, subpharyngeal branch of V_2/3A_; rV_2/3B_, third branch of trigeminal nerve; ulm, upper lip muscle; ulp, upper lip; vel, velum; vm, velar muscle. Scale bars: 100 µm for (A, applied also for B; I, applied also for J) and 50 µm for (C, applied for C–H; K, applied also for L).

## Discussion

Previous anatomical and developmental studies have demonstrated that the trigeminal nerve of the lamprey comprises three main branches in a similar manner to that of the gnathostomes (Gilland and Baker, 2005; Johnston, 1905; Koyama et al., 1987; Kuratani et al., 1997; Lindström, 1949). Still, there is a significant difference between the lamprey trigeminal nerve and that of the extant jawed vertebrates; the second branch of the former (i.e., the rV_2/3A_) contains motor components, while the second branch of the latter (i.e., the rV_2_) lacks them with an exception of holocephalans (Edgeworth, 1935; Mallatt, 1996, 2008). Furthermore, it has been shown that the rV_2/3A_ of the lamprey has a sub-branch (i.e., the rV_2/3Asubp_) that extends to the lower lip, suggesting that the lamprey mouth is controlled predominantly by rV_2/3A_ and nott, or at least less significantly, by rV_2/3B_, which has been generally regarded as homologous to the rV_3_ of the jawed vertebrates (Gaskell, 1900). These differences imply a possibility that a drastic modification of mouth control had occurred during the evolutionary transition from jawless to jawed vertebrates. Nevertheless, it has not been confirmed whether the rV_2/3Asubp_ actually contains motor fibers (Gaskell, 1900) or it is just somatosensory (Johnston, 1905).

To address this issue, we conducted whole-mount AChE staining for the lamprey prolarvae to visualize cholinergic neural tissues, followed by gene expression and histological analyses to distinguish motor fibers from motoneuronal somata and muscles.

**Motor component in the rV**_2/3Asubp_

In this study, we demonstrated that AChE-staining signals were present in the upper lip, the eminence of the buccal cavity, the medial velum region, and the pVMLB. These regions are innervated by the trigeminal nerve (Damas, 1944; Johnston, 1905; Lindström, 1949; Tretjakoff, 1929), diving into three main branches: the ophthalmic nerve (the rV_1_), the rV_2/3A_, and the rV_2/3B_. The rV_1_ sensorily innervates the dorsal part of the upper lip and contains no motor component, consistent with our results of the AChE staining. The sub-branches of the rV_2/3A_ and rV_2/3B_ have been well described in previous studies. For example, Johnston (1905) and Lindström (1949) described V_2/3A_ as r. subopticus, which ramifies into r. basilaris and r. apicalis. Subsequently, r. subpharyngeus branches (rV_2/3Asubp_) from r. basilaris. In addition, Johnston (1905) reported that r. apicalis, r. basilaris, and r. mandibularis (rV_2/3B_) contain motor components. However, these authors regarded that the rV_2/3Asubp_ are only somatosensory.

In the present study, we found the AChE staining signal in the nerve fibers passing through the pVMLB. Yokoyama et al. (2021) described two cranial nerve bundles in the subpharyngeal region ventral to the velum: rV_2/3B_ and a sub-branch of the rV_2/3A_ (i.e., the rV_2/3Asubp_). They also showed that the rV_2/3B_ and rV_2/3Asubp_ bundles distribute in lateral and medial parts, respectively. As the AChE staining signal we found in the ventral part of the pVMLB corresponds to the medial part, our results strongly suggest that rV_2/3Asubp_ contains motor components, controlling the lower lip muscles.

### The trigeminal nerve branches of the larval lamprey mark the boundary between the pre-mandibular and the mandibular regions, providing an insight into the evolutionary origin of the vertebrate jaw

The anterior ventral part of the vertebrate head is generally regarded as consisting of two developmental modules: the pre-mandibular (pre-MA) and the mandibular (MA) domains (Couly et al., 1993; Kimmel and Eberhart, 2008; Wada et al., 2011). A previous study suggested that the lamprey upper and lower lip are innervated by rV_2/3A_ and rV_2/3B_, belonging to the pre-MA and MA domains, respectively (Kuratani et al. 2004).

However, the present study sheds light on another possibility that the lower lip is controlled by the rV_2/3A_ and thus a part of the pre-MA. Consistently, Yokoyama et al. (2021) showed that the upper and lower lip receive the pre-MA stream of the neural crest cells, distinct from the posterior one migrating to the MA or velar region. Therefore, the oral apparatus-innervating rV_2/3A_ and the velum-innervating rV_2/3B_ appears to mark the boundary between the pre-MA and MA domains.

This interpretation leads to a hypothesis that the lamprey rV_2/3A_ is homologous to the rV_3_ of the jawed vertebrates (the “rV_2/3A_ = rV_3_” hypothesis), contrary to the common assumption that the lamprey rV_2/3B_ is the homolog of the rV_3_ of the jawed vertebrates (the “rV_2/3B_ = rV_3_” hypothesis). As another line of evidence for the “rV_2/3A_ = rV_3_” hypothesis, Barreiro-Iglesias et al. (2011) revealed that doublecortin (DCX), a marker of immature or migrating neurons, is expressed not in upper and lower lips-innervating (i.e., rV_2/3A_) motoneurons but in velum-innervating (i.e., rV_2/3B_) ones. Since the lower jaw-innervating motoneurons of rV_3_ in jawed vertebrates are also DCX-negative (Capes-Davis et al., 2005; Gleeson et al., 1999; Kim et al., 2006; Reiner et al., 2006; Walker et al., 2007), this observation suggests that the both the lamprey rV_2/3A_ and the rV_3_ of the jawed vertebrates share the homologous neuronal subtype. Furthermore, Tamura et al. (2023) showed that the sensory neurons express *Hmx1* in the lamprey rV_2/3A_ and the rV_3_ of the jawed vertebrates, further supporting the “rV_2/3A_ = rV_3_” hypothesis.

If it is the case that the “rV_2/3A_ = rV_3_” hypothesis is true, then is there any trigeminal nerve branch that corresponds to the lamprey rV_2/3B_ in jawed vertebrates? Probably not, because there is no known organ homologous to the lamprey velum in extant jawed vertebrates. Nevertheless, fossil “jaw-less gnathostomes” may have possessed both mouth-manipulating muscles and the velar organ innervated by different rV_2/3_ branches (Janvier, 1985, 1996; Wängsjö, 1952). The acquisition of the jaw probably caused a drastic reorganization of the anterior craniofacial region, including a heterotopic shift of oral (i.e., pre-mandibular) developmental mechanisms to the mandibular arch (Kuratani, 2005, 2012; Kuratani et al., 2004; Shigetani et al., 2005; Shigetani et al., 2002) and a loss of ancestral rV_2/3B_-equivalent branch along with velum-like structure(s) under its motor control (Fig. 6).

**Fig. 6.**
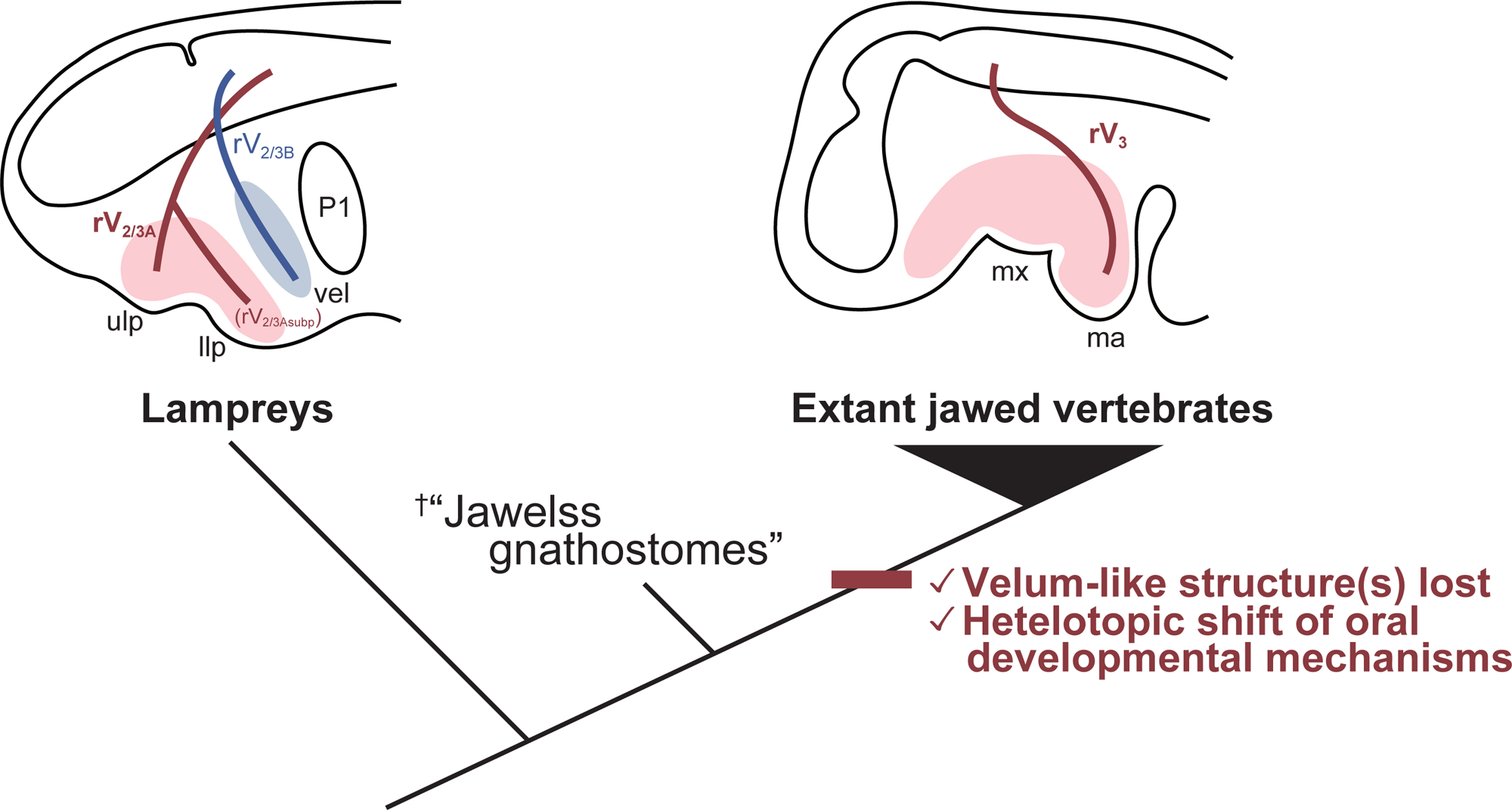
Schematic diagram for the evolution of the jaw, associated with that of the trigeminal motor branches. Motor components of the trigeminal nerve branches are visualized. Abbreviations: llp, lower lip; ma, mandible; mx, maxilla; P1, first pharyngeal pouch; rV_2/3A_, second branch of the lamprey trigeminal nerve; rV_2/3Asubp_, subpharyngeal branch of V_2/3A_; rV_2/3B_, third branch of the lamprey trigeminal nerve; rV_3_, third branch of the gnathostome trigeminal nerve; uj, upper jaw; ulp, upper lip; vel, velum.

## Author Contributions

M.T. and D.G.S. conceived the project, designed experiments, performed experiments and wrote the manuscript.

## Acknowledgments

We would like to express our thanks to Mr. Tadashige Kishi for providing live specimens of animals. The present study was supported in part by JSPS KAKENHI Grant Numbers JP22K15164 and JP24K09556 (to D.G. Suzuki).

## Conflicts of Interest

Not applicable.

## Notes

### Competing Interest Statement

The authors have declared no competing interest.

