## Supplement for "Whole-mount acetylcholinesterase (AChE) staining reveals unique motor innervation of the lamprey oral region: with special reference to the evolutionary origin of the vertebrate jaw"

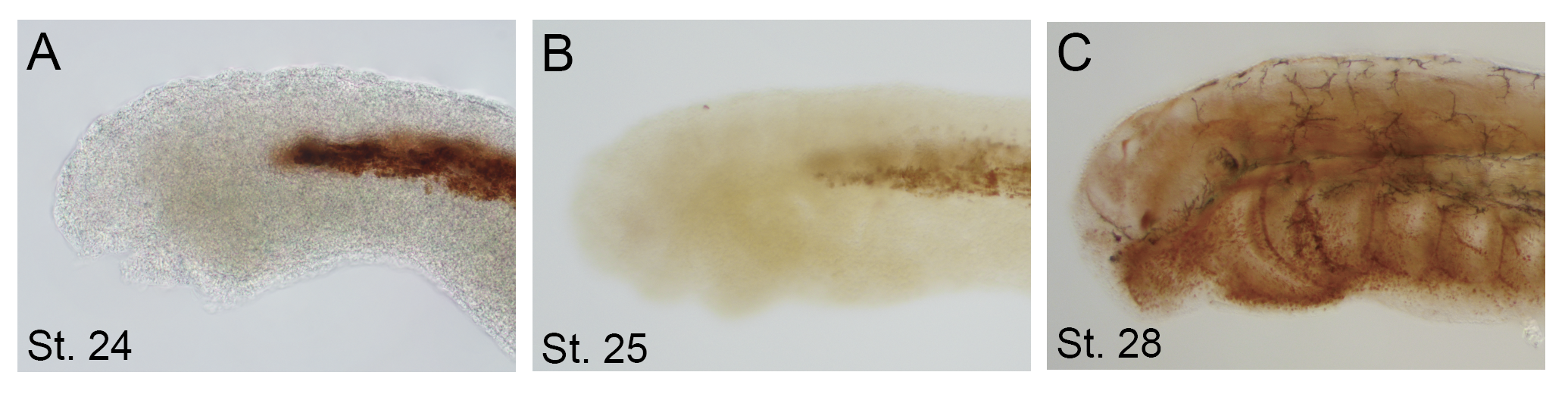

**Figure S1. Whole-mount AChE staining of lamprey embryos and prolarva.** (A) St. 24, (B) St. 25, (C) St. 28.
